# Apical progenitors remain multipotent throughout cortical neurogenesis

**DOI:** 10.1101/478891

**Authors:** Polina Oberst, Sabine Fièvre, Natalia Baumann, Cristina Concetti, Denis Jabaudon

## Abstract

The diverse subtypes of excitatory neurons that populate the neocortex are born from progenitors located in the ventricular zone (apical progenitors, APs). During corticogenesis, APs progress through successive temporal states to sequentially generate deep- followed by superficial-layer neurons directly or *via* the generation of intermediate progenitors (IPs). Yet little is known about the plasticity of AP temporal identity and whether individual progenitor subtypes remain multipotent throughout corticogenesis. To address this question, we used FlashTag (FT), a method to pulse-label and isolate APs in the mouse neocortex with high temporal resolution to fate-map neuronal progeny following heterochronic transplantation of APs into younger embryos. We find that unlike daughter IPs, which lose the ability to generate deep layer neurons when transplanted into a younger host, APs are temporally uncommitted and become molecularly respecified to generate normally earlier-born neuron types. These results indicate that APs are multipotent cells that are able to revert their temporal identity and re-enter past molecular and neurogenic states. AP fate progression thus occurs without detectable fate restriction during the neurogenic period of corticogenesis. These findings identify unforeseen cell-type specific differences in cortical progenitor fate plasticity, which could be exploited for neuroregenerative purposes.

---

During neocorticogenesis, distinct subtypes of neurons are sequentially generated, and can be distinguished by their laminar location, connectivity, and gene expression programs (Greig et al., 2013; Jabaudon, 2017). They are born from dynamic subtypes of progenitors (*i.e*. apical and intermediate progenitors) whose molecular identities and neurogenic potential are increasingly understood (Gaspard et al., 2008; Gao et al., 2014; Okamoto et al., 2016; Yuzwa et al., 2017; Mihalas and Hevner, 2018), but whose corresponding fate potential remains unknown. While the aggregate ability of cortical progenitor populations to generate earlier-born neuron types when transplanted in a younger host appears to be lost (Frantz and McConnell, 1996; Desai and McConnell, 2000), whether individual subtypes of progenitors have the potential to revert to previous temporal states remains unexamined. Here, we take advantage of a recently developed method to isolate specific populations of cortical progenitor cells (Telley et al., 2016; Govindan et al., 2018) and probe their fate potential using heterochronic transplantation into younger hosts.

In order to interrogate the temporal plasticity of AP identity, we performed FlashTag (FT) pulse-labeling in CAG::mRFP1 donor mice and immediately isolated FT-labeled cells using flow cytometry (FT specifically labels APs based on their juxtaventricular location) (Telley et al., 2016; Govindan et al., 2018). These FT^+^ APs were transplanted into wild-type (WT) host embryos by intraventricular injection (Nagashima et al., 2014) (Fig. 1, Fig. S1a), and the fate of their (RFP^+^) neuronal progeny was determined on postnatal day (P) 7, once migration is complete. To unequivocally identify neurons born in the host, we chronically administered EdU to the host dam from the time of transplantation on (Farah, 2004), such that host-born transplanted neurons could be identified as RFP^+^ EdU^+^ cells. We first performed isochronic AP transplantations at E15.5 (AP_15→15_), a time at which superficial layer (SL) neurons (*i.e.* neurons in layers (L) 4 and 2/3) are born, and at E12.5 (AP_12→12_), a time at which deep-layer (DL) neurons (*i.e.* L6 and L5) are generated (Greig et al., 2013; Jabaudon, 2017). Using these two conditions as controls, we then heterochronically transplanted E15.5 APs into E12.5 host embryos (AP_15→12_).

**Figure 1:**
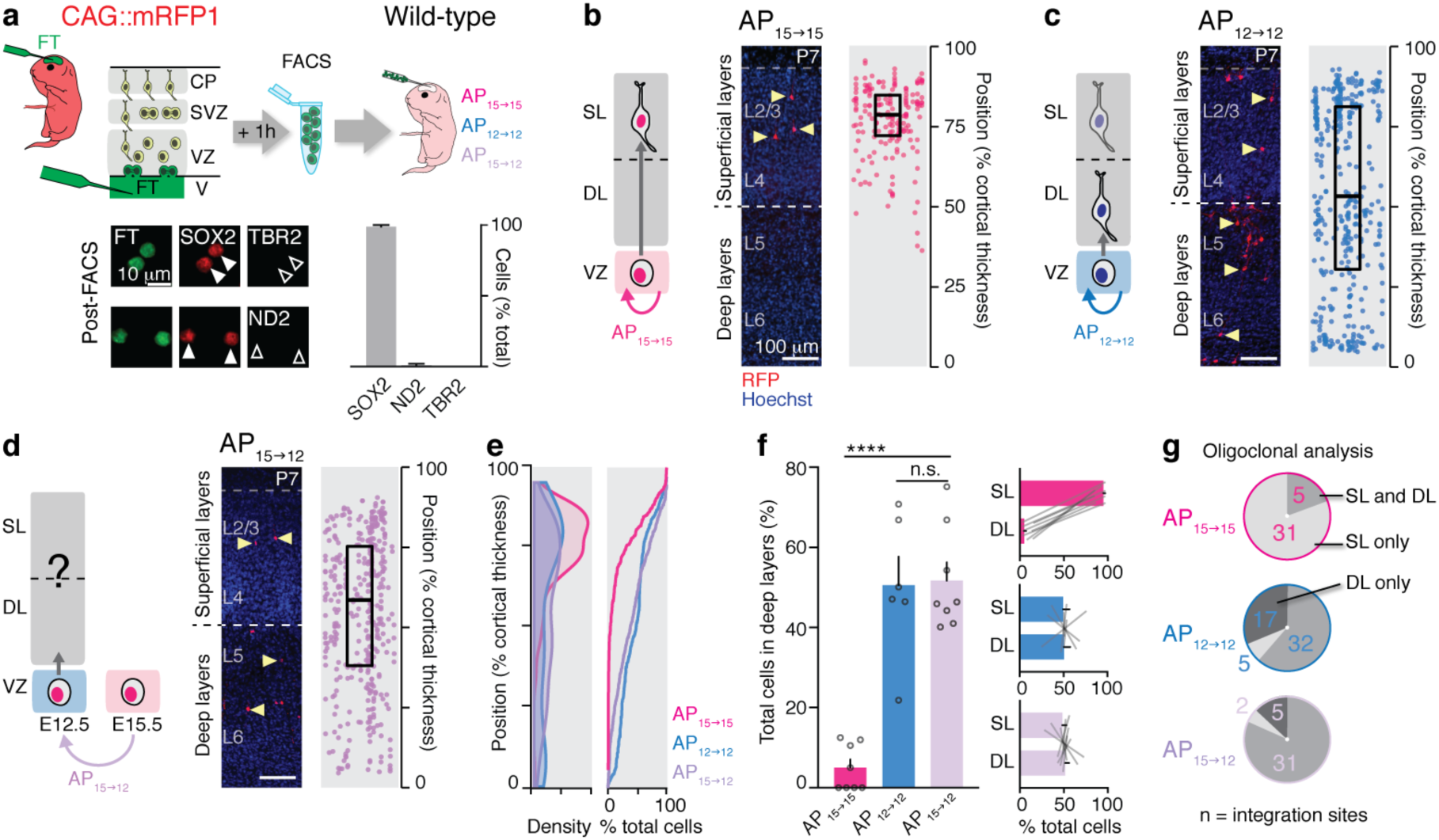
E15.5 apical progenitors (APs) remain competent to generate earlier-born deep layer neurons. **a**, Top: Schematic representation of the AP isolation and transplantation procedure. Bottom: Donor cells consist essentially of SOX2^+^ APs. TBR2: IP marker; ND2 (NeuroD2): neuronal marker. **b**, Isochronically transplanted E15.5 APs (AP_15→15_) generate superficial layer (SL) neurons. **c**, Isochronically transplanted E12.5 APs (AP_12→12_) give rise to deep layer (DL) and SL neurons. **d**, E15.5 APs transplanted into an E12.5 host (AP_15→12_) give rise to DL and SL neurons. **e**, Radial distribution of daughter neurons at P7. **f**, Left: Fraction of daughter neurons in deep layers at P7. One-way ANOVA with post-hoc Tukey test. Right: Modal distribution of daughter neurons in SL *vs*. DL. **g**, Oligoclonal integration site analysis of the laminar distribution of daughter neurons (see also Fig. S1b). CP: cortical plate; FT: FlashTag; SVZ: subventricular zone; V: Ventricle; VZ: ventricular zone. ****: P < 0.0001. b-d: on right panels, vertically-aligned cells belong to single integration sites.

Integration of donor APs into the host ventricular zone (VZ) occurred at discrete sites (typically 5-6 integration sites per host embryo, with an average of 2 donor APs per integration site, Fig. S1b). Within 24 hours, donor cells in the VZ had a typical radial glia morphology, including a radial process extending to the pial surface and juxtaventricular mitosis (Florio and Huttner, 2014; Govindan and Jabaudon, 2017) (Fig. S1c). Over the course of several days, the progeny of transplanted APs migrated away from the VZ, following a time course that was similar to that of endogenous cells (Fig. S1d). When examined at P7, both AP_15→15_ and AP_12→12_ had generated daughter neurons with appropriate laminar locations: AP_15→15_ gave rise to SL neurons (Fig. 1b, Fig. S2, Supplementary Table 1) while AP_12→12_ gave rise to DL and SL neurons, consistent with the sequential production of DL followed by SL neurons (Fig. 1c, Fig. S3, Supplementary Table 1). At each integration site, about 8 neurons were found at P7; these numbers are consistent with these cells being the progeny of mostly 1-2 progenitors (Gao et al., 2014), *i.e.* “oligoclones”, as reported above (Fig. S1b and Methods). The laminar distribution of daughter neurons in both the AP_15→15_ and AP_12→12_ conditions was replicated by *in utero* electroporation of a piggyBac-transposon construct (which allows fate mapping by genomic integration of a fluorescent protein-coding transgene into progenitors (Chen and LoTurco, 2012)), indicating that the transplantation procedure itself does not detectably affect AP neurogenic competence (Fig. S4, Supplementary Table 1).

In striking contrast to the normally exclusive SL location of AP_15→15_ daughter neurons, in the AP_15→12_ condition, daughter neurons were located in both DL and SL, as described above for AP_12→12_ (Fig. 1d-f, Fig. S5a, b, Supplementary Table 1). This laminar redistribution was also present when examining individual oligoclonal integration sites (Fig. 1g). This consistent laminar redistribution across all integration sites argues against a population based effect in which only few transplanted cells are highly plastic, but instead suggests that essentially all integrated APs are undergoing respecification. Together, these data suggest that E15.5 APs remain competent to generate DL neurons.

DL neurons in the AP_15→12_ condition could in principle reflect a mismigration of this fraction of cells, which nonetheless retain SL molecular identities. To investigate this possibility, we examined the molecular identity and axonal projections of daughter neurons (Fig. 2). Molecular identity was assessed using TBR1, a marker of L6 neurons, CTIP2, a marker of L5 neurons, and CUX1, a marker of L2-4 neurons (Greig et al., 2013). Expression of these molecular markers in AP_15→12_ daughter neurons was congruent with laminar position: CUX1 was expressed by SL neurons while CTIP2 and TBR1 were expressed by DL neurons (Fig. 2a-c, Fig. S6a-c). This harmonious laminar and molecular identity strongly suggest a respecification of the AP_15→12_ progeny. Further supporting this possibility, in contrast to the neuronal progeny of E15.5 APs, which send axons to intracortical but not to subcerebral targets (Greig et al., 2013), the AP_15→12_ progeny also projected subcerebrally, as do E12.5 AP daughter neurons (Fig. 2d). Together, these data reveal an embryonic age-appropriate and congruous re-specification of the laminar, molecular, and hodological identity of AP_15→12_ daughter neurons.

**Figure 2:**
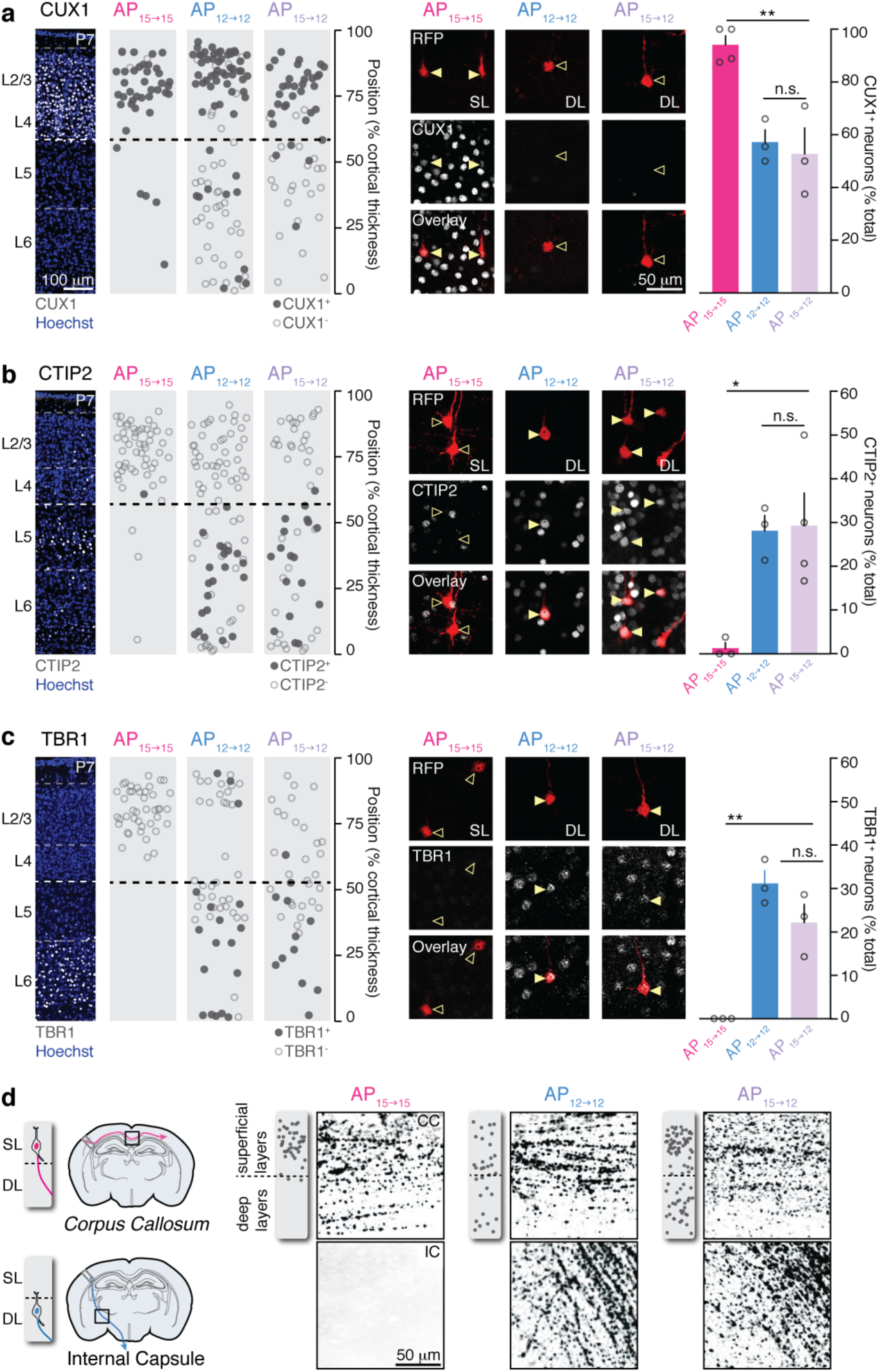
Embryonic age-appropriate molecular features and connectivity of AP_15→12_ daughter neurons. **a**, CUX1, a superficial layer (SL) neuron marker, is reduced in deep layer (DL) AP_15→12_ daughter neurons. **b**, CTIP2, a DL neuron marker, is increased in DL AP_15→12_ daughter neurons. **c**, TBR1, a DL neuron marker, is increased in DL AP_15→12_ daughter neurons. a-c: one-way ANOVA with post-hoc Tukey test. **d**, AP_15→12_ daughter neurons extend subcerebral projections *via* the internal capsule. Photomicrographs show RFP+ axons from the same pup for each condition. Insets on their left show the laminar position of neurons in the corresponding pup. CC: Corpus Callosum; IC: Internal Capsule. *: P < 0.05, **: P< 0.01.

DL neuronal identity could in principle result from a post mitotic process rather than from a pre-mitotic respecification of AP_15→12_. To investigate this possibility, we examined the fate of the small population of neurons which were being born in the donor at the time of isolation. In contrast to AP_15→12_, these neurons, identified as RFP^+^ EdU^−^ cells (*i.e.* transplanted cells which never underwent division in the host) still migrated to SL, as they would have done in their original host, indicating that they were already committed to an SL fate (Fig. S6d,e). Thus, emergence of DL neurons in the AP_15→12_ condition appears to result from a pre-mitotic rather than a post-mitotic process.

Do AP_15_ acquire an AP_12_ identity upon heterochronic transplantation? To directly identify potential changes in the molecular identity of AP_15→12_, we performed patch-seq RNA sequencing (Cadwell et al., 2016; Fuzik et al., 2016) of visually-identified RFP^+^ juxtaventricular cells (*i.e.* presumptive APs) (Fig. 3a). We first determined the embryonic age-specific transcriptional identity of normal AP_15_ and AP_12_ and compared AP_15→12_ identity to these two control conditions using a linear regression model (Teo et al., 2010). Similarly, we generated an additional model to specifically identify genes distinguishing AP_15→12_ from normal AP_15_ (*i.e.* to identify genes whose expression changes upon heterochronic transplantation of AP_15_) (Fig. 3b,c, Fig. S7). This two-pronged approach revealed that AP_15→12_ repress AP_15_-type transcriptional programs and re-induce AP_12_-type transcriptional programs, and identified dynamically regulated genes following heterochronic transplantation (Fig. 3b,c, Fig. S7). Re-induced, normally early-expressed genes included the Wnt pathway regulator *Tcf7l1*, which represses neuronal differentiation and increases self-renewal in cortical apical progenitors (Kuwahara et al., 2014) and polycomb repressive complex 2 (PRC2) methyltransferase *Ezh2*, in the absence of which APs undergo precocious neurogenesis (Pereira et al., 2010). Both of these genes thus act on the dynamically regulated balance between self-renewal and differentiation during corticogenesis (Paridaen and Huttner, 2014). Repressed, normally late-onset, genes included *Tenascin c* (*Tnc*), an extracellular glycoprotein involved in the onset of gliogenesis late in corticogenesis (Garcion et al., 2004) and *Mfge8*, coding for an EGF-like domain containing protein, which regulates quiescence in adult neural stem cells (Zhou et al., 2018). Together, these results reveal that the temporal shift in the neurogenic competence of AP_15→12_ is accompanied by a corresponding wholesale shift in their global molecular identity.

**Figure 3:**
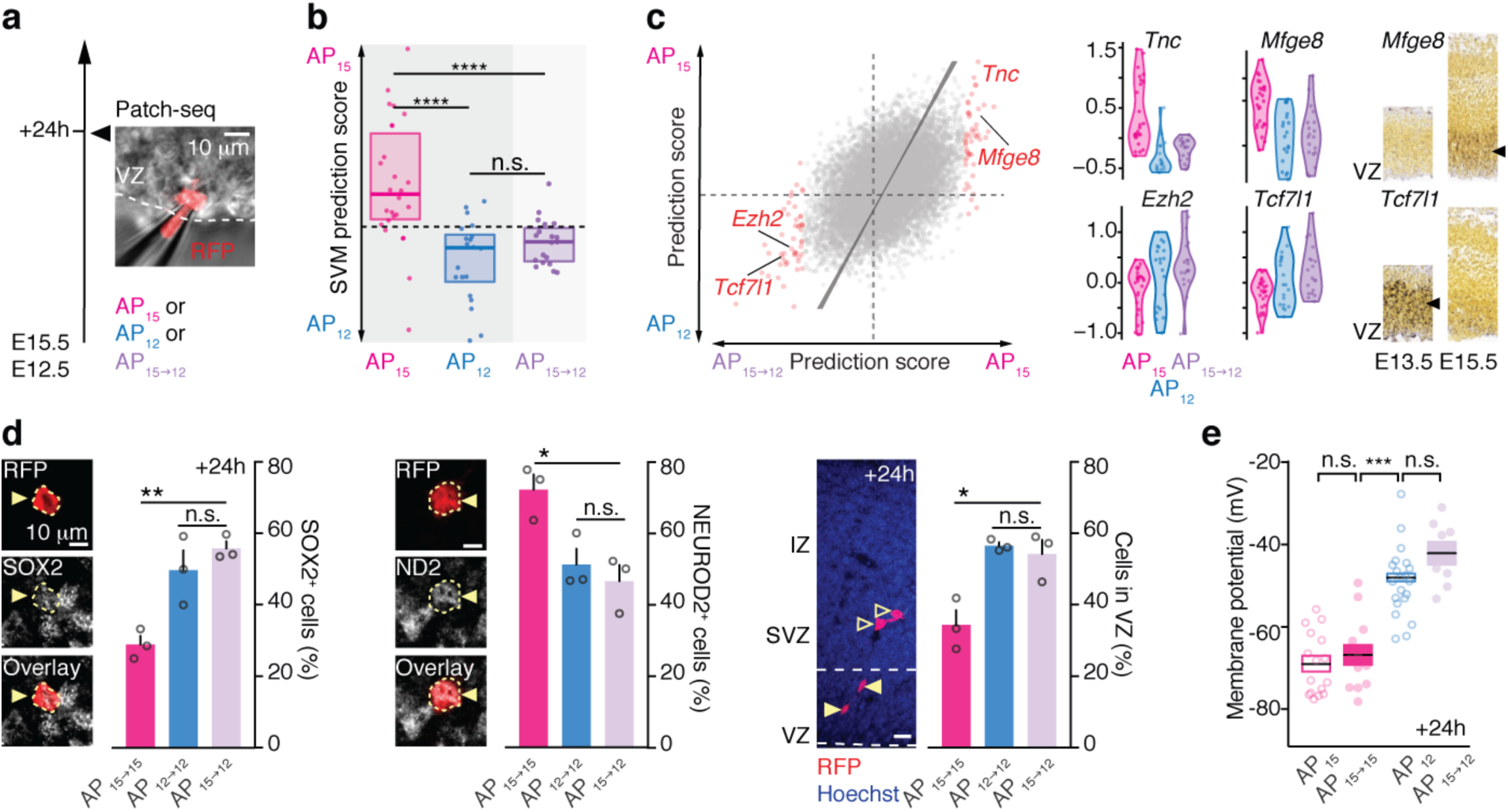
AP_15_ are respecified into AP_12_ upon transplantation. **a**, Summary of the experimental approach used for patch-seq single cell RNA sequencing. Inset: patched AP. **b**, SVM classification reveals that AP_15→12_ acquire an AP_12_-type molecular identity. Kolmogorov-Smirnov test. **c**, AP_15→12_ repress AP_15_- and induce AP_12_-enriched transcripts. Right: examples of AP_15_ and AP_12_ genes that are repressed or induced in AP_15→12_. Insets: *in situ* hybridizations from the Allen Developing Mouse Brain Atlas confirm transcriptomic data (www.brain-map.org). **d**, Neurogenic divisions decrease to AP_12_ values in AP_15→12_. Accordingly, more cells remain in the VZ. One-way ANOVA with post-hoc Tukey-test. **e**, The membrane potential of AP_15→12_ depolarizes to AP_12_ values. Kruskal-Wallis test with post-hoc Dunn’s test. IZ: intermediate zone; ND2 (NeuroD2): neuronal marker; SOX2: AP marker; SVM: support vector machine; SVZ: subventricular zone; VZ: ventricular zone. One-way ANOVA with post-hoc Tukey test. *: P < 0.05, **: P < 0.01, ***: P < 0.001, ****: P < 0.0001.

We next examined whether this E12-like molecular re-specification of AP_15→12_ was accompanied by a corresponding reassignment of their functional features (Fig. 3d,e). For this purpose, we measured two key temporally-regulated physiological parameters of these cells:

*(1) Neurogenic divisions*. Symmetric divisions predominate early in corticogenesis while asymmetric, neuron-generating divisions increase at later embryonic stages (Haubensak et al., 2004; Govindan and Jabaudon, 2017). Accordingly, RFP^+^SOX2^+^ cells (*i.e.* cycling transplanted APs) were more prevalent in the AP_12→12_ condition than the AP_15→15_ condition (Fig. 3d, left), while the reverse was true for RFP^+^NEUROD2^+^ cells (*i.e.* daughter neurons) (Fig. 3d, center). Similarly, in line with a higher self-replication rate of APs early in corticogenesis, a greater fraction of daughter cells remained in the VZ in the AP_12→12_ condition (Fig. 3d, right). Consistent with acquisition of AP_12_-like mitotic properties by AP_15→12_, SOX2^+^ daughter cells increased, NEUROD2^+^ daughter cells decreased, and daughter cells remained in the VZ, as was the case for AP_12→12_ (Fig. 3d).

*(2) Resting membrane potential*. Progression of AP neurogenic competence is regulated by a progressive hyperpolarization of the resting membrane potential (V_m_) in a Wnt-dependent manner (Vitali et al., 2018). Since the Wnt pathway regulator *Tcf7l1* was re-induced in AP_15→12_ (Fig. 3c), we hypothesized that a resetting of V_m_ values might contribute to the re-specification process. To examine this possibility, we measured V_m_ values in transplanted APs using whole-cell patch clamp recording. V_m_ values in isochronically-transplanted cells did not differ from their host counterparts, indicating that this electrophysiological parameter is not affected by the transplantation procedure. In contrast,

AP_15→12_ V_m_values were reset to those of APs in their E12.5 host, revealing a reassignment of this critical regulator of AP competence, potentially contributing to neurogenic re-specification (Fig. 3e).

Together, the results above reveal a reset in the molecular and physiological properties of AP_15→12_, which are reassigned to those of their younger host. Our results contrast with earlier findings in which progenitor cells transplanted into younger hosts could no longer generate DL neurons (Frantz and McConnell, 1996; Desai and McConnell, 2000). In these seminal experiments, progenitors were identified by incorporation of radiolabeled thymidine, which labels S-phase cells across the VZ, subventricular zone and outer subventricular zone (Angevine and Sidman, 1961). Late in corticogenesis, and particularly in the ferret (the species in which these experiments were performed), the majority of progenitors are IPs (Reillo and Borrell, 2012). In these studies, the fate plasticity identified here may thus have been occluded by the overwhelming predominance of other, potentially less plastic, progenitor cell types, and particularly by IPs. In addition, in contrast to APs, which divide multiple times, IPs are thought to divide only once or twice before reaching a terminal division (Mihalas et al., 2016; Mihalas and Hevner, 2018). As a consequence, radiolabeled thymidine, which was used in these early studies, will be more diluted in the neuronal progeny of APs than in the progeny of IPs, potentially biasing quantifications towards the latter lineage.

To investigate the possibility of cell-type specific differences in progenitor plasticity, we first reproduced the strategy used to isolate progenitors in these earlier experiments: following microdissection and dissociation of the dorsal pallium at E15.5, we used the thymidine analog EdU to single pulse-label VZ and SVZ progenitors and identify their progeny in the host (Fig. 4a). Following transplantation into E12.5 hosts, daughter neurons, identified as EdU^+^ cells, were mostly located in SL, replicating earlier findings in the ferret (Fig. 4b, Fig. S8, Supplementary Table 1). Thus, compared with previous studies, our new results reflect cell-type specific rather than species-specific features of progenitor plasticity, and suggest that the competence for progenitor re-specification is subtype dependent.

**Figure 4:**
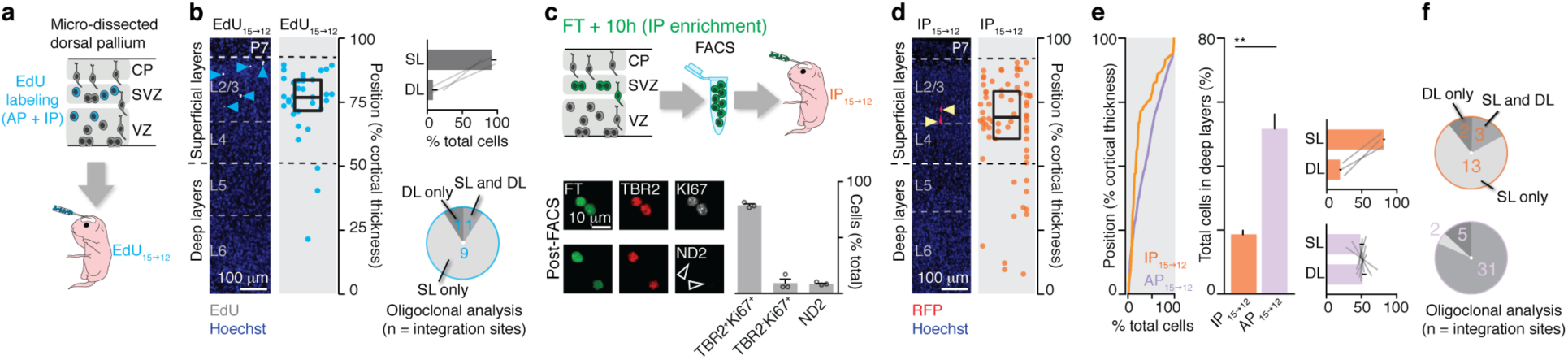
E15.5 intermediate progenitors (IPs) are committed to generating superficial layer neurons. **a**, Left: Schematic representation showing transplantation of EdU-based labeling and transplantation procedure. **b**, E15.5 EdU+ cells transplanted into an E12.5 host (EdU_15→12_) still give rise to SL neurons. Compare with Fig. 1d. Right, top: Modal distribution of daughter neurons in SL *vs*. DL. Right, bottom: Single integration site analysis of the laminar distribution of daughter neurons. **c**, Schematic representation of the IP isolation and transplantation procedure. **d**, E15.5 IPs transplanted into an E12.5 host (IP_15→12_) essentially give rise to SL neurons. **e**, Left: Radial distribution of the daughter neurons of IP_15→12_ and AP_15→12_ at P7. Center: Fraction of daughter neurons in DL at P7. Student’s t-test. Right: Modal distribution of daughter neurons in SL *vs*. DL in the IP_15→12_ condition and AP_15→12_ condition. **f**, Oligoclonal integration site analysis of the laminar distribution of daughter neurons. AP_15→12_ data in e-f have been reproduced from Fig. 1e-g for direct comparison with IP_15→12_. CP: cortical plate; FT: FlashTag; ND2 (NeuroD2): neuronal marker; SVZ: subventricular zone; TBR2: IP marker; V: Ventricle; VZ: ventricular zone. b,d: Vertically-aligned cells belong to single integration sites.

In light of these results, we hypothesized that while E15.5 APs are temporally plastic, E15.5 daughter IPs instead are committed to generating SL neurons. To test this possibility, we performed FT labeling at E15.5 and waited 10 hours before collection to allow daughter IPs to differentiate (Telley et al., 2016; Govindan et al., 2018). At this stage, > 75% of FT^+^ cells have differentiated into IPs, as revealed by co-expression of TBR2 and the proliferation marker KI67 (Vitali et al., 2018) (Fig. 4c, Fig. S9a). Upon examination of single integration sites at P7, transplanted IPs gave rise to a smaller number of cells than APs, consistent with a more differentiated status and a limited number of divisions (Gao et al., 2014; Mihalas et al., 2016; Mihalas and Hevner, 2018) (Fig. S9b). In contrast to AP_15→12_, IP_15→12_ essentially gave rise to SL neurons (Fig. 4d-f, Fig. S9c, Supplementary Table 1). Consistent with their location, these neurons expressed CUX1 but not CTIP2 (Fig. S9d). Similarly, as predicted by the lack of temporal plasticity of newborn neurons (Fig. S6e) 10 hour-old E15.5 neurons (identified as RFP^+^EdU^−^ cells, *i.e.* transplanted cells which never underwent division in the chronic EdU-perfused host, see Fig. S6d) did not change their fate when transplanted into E12.5 hosts (Fig. S9e). These results demonstrate that in contrast to their mother cells, differentiated IPs are committed progenitors with a fixed neurogenic competence.

Together, our findings reveal an outstanding ability of APs to revert their temporal identity and re-enter past molecular states to once again generate normally earlier-born neurons. In contrast, IPs are committed progenitors which lack such fate plasticity. These results highlight an unexpected cell-type specific diversity in the temporal plasticity of neocortical progenitors. Moreover, they reveal that AP fate progression occurs without detectable fate restriction during the neurogenic period of corticogenesis. Thus, although the arc of development in whole organisms inescapably extends towards maturity, subsets of cells can be untethered from this course and remain able to re-enter past developmental states.

While technical limitations had so far precluded access to the cell-type specific temporal plasticity identified here, the ability of progenitor cells to at least partially revert to earlier embryonic states has been described in other settings. For example, in *Drosophila* neuroblasts, late re-expression of early temporal identity factors leads to the production of previously generated cells (Cleary and Doe, 2006; Kohwi and Doe, 2013; Kohwi et al., 2013). Similarly, re-expression of the early-neuron transcription factor *Fezf2* at later corticogenic stages leads to re-emergence of neurons with deep-layer identities (Molyneaux et al., 2005), and inactivation of the transcription factor *Foxg1* late in development leads to resurgence of early-born Cajal-Retzius neurons (Hanashima, 2004). Together with classical cell biology work showing that nuclear transfer from adult somatic cells into enucleated oocytes leads to nuclear reprogramming to younger states (Gurdon, 1962; Wakayama et al., 1998), these examples indicate that the transcriptional programs used during development remain amenable to re-recruitment even after progenitor fate has progressed.

The temporal resetting in V_m_ values identified here may contribute to the reassignment in AP identity and regulate sensitivity to extracellular signals (Vitali et al., 2018). Such temporally dynamic environmental factors may include cell-cell interactions (Mutch et al., 2009; Okamoto et al., 2016), metabolic supplies, neurotransmitter-mediated feedback from newborn neurons or nearby axons (Seuntjens et al., 2009; Toma et al., 2014), as well as the protein and ion composition of the neurogenic niche and cerebrospinal fluid (Lehtinen et al., 2011), which together may be involved in AP re-specification. It will be important, in future studies, to understand the precise sequence of extrinsic and intrinsic, permissive and instructive events underlying such a remarkable level of plasticity, and attempt to harness these processes for neuroregenerative purposes.

## Supporting information

## Acknowledgements

We thank Eiman Azim and members of the Jabaudon lab for comments on the manuscript. We thank Olivier Raineteau for providing plasmids. We thank Andrea S. Lopes, Audrey Benoit, the FACS facility, the Genomics Platform and the Bioimaging Facility of the University of Geneva for technical assistance. Work in the Jabaudon laboratory is supported by the Swiss National Science Foundation. P.O. is supported by the iGE3.

## Author Contributions

P.O. performed the experiments with the help of C.C.; S.F. performed the electrophysiology experiments and the collection of cells for patch-seq with the help of P.O.; N.B. performed the bioinformatic analysis; P.O. and D.J. wrote the manuscript.

## Data and material availability

Annotated data and accession numbers will be available upon manuscript publication. The supplementary materials contain additional data.

## Methods

### Mice

Experiments were performed using CD1 (Charles River) and CAG::mRFP1 (B6.Cg-Tg(CAG-mRFP1)1F1Hadj/J, JAX 005884) (Long et al., 2005) mice. CAG::mRFP1 mice were bred on a CD1 background for at least 10 generations before use. Embryonic day (E) 0.5 was established as the day of vaginal plug. Both female and male embryos were used throughout the study. All experimental procedures were approved by the Geneva Cantonal Veterinary Authorities, Switzerland.

### *In utero* FlashTag labeling

FT injections were performed at E12.5 and E15.5 as previously described (Govindan et al., 2018; Telley et al., 2016). Briefly, pregnant mice were anaesthetized with isoflurane and an abdominal incision was made to access the uterine horns. Half a microliter of 10 mM carboxyfluorescein succinimidyl ester (“FlashTag”; CellTraceTM CFSE, Life Technologies, #C34554) was injected into the lateral ventricles of embryos using a picospritzer. The abdominal wall was sutured and the embryos let to develop until collection.

### Preparation of cell suspension and *in utero* cell transplantation

One hour after FT labeling (unless otherwise stated) E12.5 or E15.5 donor embryos were collected in ice-cold HBSS and the dorsal pallium was microdissected under stereomicroscopic guidance (Leica, #M165 FC) using a microscalpel. Tissue from 6-8 littermates was pooled and incubated in 400 µl TrypLE (Gibco, #12605-010) for 3 minutes at 37 °C. TrypLE was inactivated by adding HBSS containing 0.1% BSA and the tissue was mechanically dissociated by triturating with a 1 ml pipette. Cells were filtered through a 70 µm cell strainer, centrifuged (150 rpm, 5 minutes) and resuspended in FACS buffer (L15 medium containing 2 mg/ml glucose, 0.1% BSA, 1:50 citrate phosphate dextrose, 10 units/ml DNase I and 1 µM MgCl_2_). Cells were sorted on an S3e Cell Sorter (BioRad) or a BD FACS Aria II flow cytometer (BD Biosciences) and the top 5 – 10% brightest cells (Govindan et al., 2018; Telley et al., 2016), were collected in ice-cold FACS buffer. Sorted cells were centrifuged (150 rpm, 5 minutes) and resuspended in HBSS containing 10 mM EGTA (Nagashima et al., 2014) and 0.1% fast green to a final density of 25,000 – 75,000 cells per µl. Cells were kept on ice until the start of the transplantation procedure. Five hundred µl of the cell suspension (corresponding to approximately 12,500 – 37,500 cells) was injected into the lateral ventricles of each host embryo using a picospritzer.

### Continuous EdU labeling

A 10 mg/ml EdU solution was prepared in (1:1) DMSO:water. An osmotic pump (Alzet, #1003D) was filled with the solution and pre-incubated at 37 °C for 4 - 12 hours. The pump was placed in the peritoneal cavity of the mother at the end of the surgical procedure to allow for continuous administration of EdU. If continuous delivery was required for more than 3 days (*i.e.* when started at E12.5), a new pump prepared in the same way as above was introduced after 72 hours. Revelation of EdU was performed using Click-it chemistry following the manufacturer’s instructions (Invitrogen).

### Oligoclonal analysis

Integration sites were defined as sites containing transplanted cells that are separated from other transplanted cells by at least 500 μm in either direction. Cells within each integration site correspond to progeny of a few integrated cells (see Fig. S1b).

### *In utero* electroporation

*In utero* electroporations were performed as previously described (Vitali et al., 2018). Briefly, a 1.5 µg/µl DNA solution containing 1:2 transposon pPB-CAG-EmGFP and hyperactive piggyBac transposase (pRP-Puro-CAGG-Pbase) (Chen and LoTurco, 2012) was prepared in sterile PBS containing 0.01 mg/mL fast green. Half a microliter of DNA solution was injected unilaterally into the lateral ventricle of embryos using a picospritzer and electroporation was performed by applying 5 electric pulses (25 V for E12.5, 45 V for E15.5; 50 ms at 1 Hz) with a square-wave electroporator (Nepa Gene, Sonidel Limited, UK). Embryos were let to develop until collection at P7.

### Immunohistochemistry

Embryonic brains were collected in PBS and fixed in 4% paraformaldehyde (PFA) overnight at 4°C. Postnatal mice were perfused with 4% PFA and brains were post-fixed in 4% PFA overnight at 4°C. Fifty to 100 µm coronal sections were cut using a vibrating microtome (Leica, #VT100S). Antigen-retrieval was performed by incubating the sections in citrate buffer solution (Dako, #S1699) for 20 minutes at 82°C. Sections were rinsed three times in PBS and incubated overnight at 4°C with primary antibodies diluted in PBS containing 3% BSA and 0.3% Triton-X-100. Sections were rinsed three times in PBS containing 0.1% Tween20 and incubated for 1 hour at room temperature with corresponding secondary antibodies (1:500, Life Technologies). Sections were washed twice with PBST and incubated with Hoechst for 5 minutes (1:5000 in PBS, Life Technologies, #33342) to stain nuclei. Sections were dry mounted on slides using Fluoromount (Sigma, # F4680).

### Immunocytochemistry of FAC-sorted cells

Sorted cells were resuspended in FACS buffer at a density of 25,000 cells per ml, seeded on glass coverslips coated with poly-d-lysine and laminin and maintained at 37 °C with 5% CO_2_ for 30 minutes to allow cells to adhere to the coverslips. Cells were fixed with 4% PFA (15 min at RT), rinsed three times with PBS and permeabilized by incubation with 0.25 % Triton-X-100 in PBS for 10 minutes, followed by three washes in PBS. Blocking was performed with 1% BSA in PBST for 1 hour, followed by incubation with primary antibodies for 2 hours at room temperature. Cells were rinsed three times in PBS and incubated for 1 hour at room temperature with corresponding secondary antibodies (1:500, Life Technologies). Sections were washed two times with PBS and incubated with Hoechst for 5 minutes (1:5000 in PBS) to stain nuclei. Coverslips were mounted on slides using Fluoromount.

### Antibodies

Rat anti-CTIP2 (1:500; Abcam, #AB18465); rabbit anti-CUX1 (1:500; Santa Cruz; sc-13024); rabbit anti-dsRed (1:1000; Clonetech; # 632496); rabbit anti-KI67 (1:250; Abcam, #AB15580); rabbit anti-NEUROD2 (1:1000; Abcam, #AB104430); rat anti-RFP (1:500; Chromotek; # 5F8); goat anti-SOX2 (1:500; SC Biotech, #SC17320); mouse anti-SOX2 (1:500; Santa Cruz; sc-365823); rabbit anti-TBR1 (1:500; Abcam; # AB31940); rabbit anti-TBR2 (1:500; Abcam; ab23345); rat anti-TBR2 (1:500; Invitrogen, #14-4875-82).

### Image acquisition, quantification and statistical analyses

Images were acquired using an Eclipse 90i epifluorescence microscope (Nikon), a LSM 700 line scan confocal (Carl Zeiss) or a Nikon A1r spectral line scan confocal (Nikon) and analyzed with ImageJ software. For analysis of transplantations, brains from each recipient litter (*i.e.* containing cells from one independent FACS experiment/one donor litter) were pooled for analysis and considered as one replicate (“N”), except when indicated otherwise. Only neurons located in the neocortex were included in analysis, and glia were excluded based on morphology. All results are shown as mean ± SEM, except when indicated otherwise. The following convention was used: *: *P <* 0.05, **: *P <* 0.01, ***: *P <* 0.001, ****: *P <* 0.0001.

**Figure 1a and 4c:** Three to four coverslips (= N) from 2 independent FACS experiments were analyzed for each condition. Figure 1a: 73 cells were stained for SOX2 and ND2 (% SOX2^+^: 98.91 ± 1.09; % ND2^+^: 1.09 ± 1.09); 50 cells were stained for TBR2 (% TBR2^+^: 0). Figure 4c: 121 cells were stained for TBR2 and Ki67 (% TBR2^+^ Ki67^+^: 78.84 ± 1.41; % TBR2^−^ Ki67^+^: 9.21 ± 3.37); 214 cells were stained for ND2 (% ND2^+^: 8.4 ± 0.75).

**Figure 1b-g:** Cells from all transplantation experiments collected at P7 were included in analysis. AP_15→15_: 238 cells from 8 independent experiments (= N) and 36 integration sites; AP_12→12_: 385 cells from 6 independent experiments and 54 integration sites; AP_15→12_: 381 cells from 8 independent experiments and 38 integration sites. Figures 1b-d: The radial position of cells within the neocortex was measured and normalized to the cortical thickness. The normalized radial position for each cell was plotted and cells were aligned on the X-axis per integration site. Chronic EdU labeling was used to identify transplanted host-born neurons in all experiments, except donor litters: AP_15→15_ 1, 2, 7, 8; AP_12→12_ 2, 3, 5; AP_15→12_ 1, 3, 4, 5. Figure 1e: Density and cumulative distribution plots were used to additionally display the radial positions of cells. Figure 1f: Left: The percentage of total cells in DL per donor litter is plotted (AP_15→15_: 5.0 ± 2.05; AP_12→12_: 50.4 ± 7.11; AP_15→12_: 51.62 ± 4.53). A one-way ANOVA with post-hoc Tukey test was used. Right: The percentage of total cells in DL and SL per donor litter is plotted. SL DL values from the same experiment are connected with lines. Figure 1g: Individual integration sites were examined for each condition and the number of oligoclones containing cells in only SL, only DL or both SL and DL were plotted.

**Figure 2:** A subset of cells depicted in Fig. 1 was analyzed for molecular marker expression. The normalized radial position of labeled and non-labeled cells and the total percentage of cells expressing the respective marker were plotted. AP_15→15_: 42 cells from 4 independent experiments were stained for CUX1 (% CUX1^+^: 94.1 ± 3.42); 54 cells from 3 independent experiments were stained for CTIP2 (% CTIP2^+^: 1.28 ± 1.28); 48 cells from 3 independent experiments were stained for TBR1 (% TBR1^+^: 0). AP_12→12_: 102 cells from 3 independent experiments were stained for CUX1 (% CUX1^+^: 57.19 ± 4.69); 53 cells from 3 independent experiments were stained for CTIP2 (% CTIP2^+^: 28.13 ± 3.52); 55 cells from 3 independent experiments were stained for TBR1 (% TBR1^+^: 31.11 ± 2.94). AP_15→12_: 59 cells from 3 independent experiments were stained for CUX1 (% CUX1^+^: 52.82 ± 9.76); 81 cells from 4 independent experiments were stained for CTIP2 (% CTIP2^+^: 29.34 ± 7.43); 38 cells from 3 independent experiments were stained for TBR1 (% TBR1^+^: 22.13 ± 4.18). A one-way ANOVA with post-hoc Tukey test was used.

**Figure 3b:** A Kolmogorov-Smirnov test was used. AP_12_: 20 cells, AP_15_: 26 cells, AP_15→12_: 29 cells: 19 cells.

**Figure 3d:** Left: AP_15→15_: 56 cells from 3 independent experiments (= N) were stained for SOX2 (% SOX2^+^: 28.97 ± 2.41); 54 cells from 3 independent experiments were stained for ND2 (% ND2^+^:72.59 ± 3.88). AP_12→12_: 49 cells from 3 independent experiments were stained for SOX2 (% SOX2^+^: •± 5.57); 72 cells from 3 independent experiments were stained for ND2 (% ND2^+^: 51.24 ± 4.38). AP_15→12_: 134 cells from 3 independent experiments were stained for SOX2 (% SOX2^+^: 55.83 ± 1.97); 63 cells from 3 independent experiments were stained for ND2 (% ND2^+^: 46.56 ± 4.57). Right: For analysis of cells remaining in the VZ 24h after transplantation, cells from 3 independent experiments (= N), including a subset of cells used for SOX2/ND2 expression analysis shown in Fig. 3d, were analyzed. AP_15→15_: 120 cells total (% cells in VZ: 34.28 ± 4.13). AP_12→12_: 66 cells total (% cells in VZ: 56.6 ± 0.95). AP_15→12_: 134 cells total (% cells in VZ: 54.15 ± 3.94). A one-way ANOVA with post-hoc Dunnett test was used.

**Figure 3e:** The resting membrane potential of transplanted or non-transplanted FT labeled control cells was recorded 24 hours after transplantation or FT labeling. Recordings from at least 3 independent experiments per condition were pooled for analysis. AP_15_: −69.01 ± 1.89 mV; N = 16 cells. AP_15→15_: −66.8 ± 2.58 mV; 12 cells. AP_12_: −48.96 ± 1.82 mV; 21 cells. AP_15→12_: −42.08 ± 2.82 mV; 8 cells. A Kruskal-Wallis test with a post-hoc Dunn’s test was used.

**Figure 4b:** 34 cells from 3 independent experiments (= N) and 11 integration sites were analyzed. Left: The normalized radial position of EdU^+^ cells was plotted and cells were aligned on the X-axis per integration site. Right, top: The layer position for each cell was plotted. SL and DL values from the same experiment are connected with lines. Right, bottom: Individual integration sites were examined and the number of oligoclones containing cells in only SL, only DL or both SL and DL were plotted.

**Figure 4d-f:** 81 cells from 3 independent experiments (= N) and 18 integration sites were analyzed. For comparison, AP_15→12_ data reproduced from Fig. 1f are shown. Figure 4d: The normalized radial position of RFP^+^EdU^+^ cells was plotted and cells were aligned on the X-axis per integration site. Figure 4e: Left: Density and cumulative distribution plots were used to additionally display the radial positions of cells. Center: The percentage of total cells in DL per donor litter is plotted (IP_15→12_: 18.98 ± 1.27). A student’s t-test was used. Right: The percentage of total cells in DL and SL per donor litter is plotted. SL and DL values from the same experiment are connected with lines. Figure 4f: Individual integration sites were examined for each condition and the number of oligoclones containing cells in only SL, only DL or both SL and DL were plotted.

**Figure S1a:** The number of integrated cells per integration site was analyzed 6 hours after transplantation of AP_15→15_. A total of 93 cells from 3 independent experiments (= N), corresponding to 43 integration sites, were analyzed. Number of cells per integration site: 1-6; mean: 2.16; median: 2.

**Figure S1c:** Brains were collected at consecutive time-points after AP_15→15_ transplantation or E15.5 FT labeling. The distance of cells to the lateral ventricle was measured using ImageJ. RFP: 6h: 46 cells, 14.89 ± 1.99 μm; 12h: 21 cells, 29.54 ± 7.35; 24h: 30 cells, 68.95 ± 13.25; 48h: 30 cells, 259.4 ± 26.85; 72h: 636.2 ± 63.42. FT: 6h: 36 cells, 35.29 ± 3.78; 12h: 25 cells, 69.9 ± 5.27; 24h: 92.79 ± 8.36; 48h: 39 cells, 225.3 ± 23.97; 72h: 681.2 ± 36.43.

**Figures S2, S3, S5a, S8, S9c, S9e bottom:** Images from all cells used for analysis are shown for each condition. In the plots, the normalized radial position is shown. Cells are aligned on the X-axis per integration site. Figures S2, S3, S5: Chronic EdU labeling was used to identify transplanted host-born neurons in all experiments, except: AP_15→15_: donor litters 1, 2, 7, 8; AP_12→12_: donor litters 2, 3, 5; AP_15→12_: donor litters 1, 3, 4, 5. Figure S8: only EdU^+^ cells were included in analysis. Figure S9c: Only EdU^+^ RFP^+^ neurons were included in analysis. Figure S9e, bottom: Only EdU^−^ RFP^+^ neurons were included in analysis.

**Figure S4:** The radial position of GFP^+^ neurons within the neocortex was normalized to the cortical thickness. At least 3 pups (= N), and 3 sections from different rostro-caudal levels per pup, were used per condition. The layer position for each cell was plotted. In cortical areas with no anatomically distinguishable L4, the lowest 1/3 of L2/3 was considered as L4. Cells in L2/3 (%): PB_15_: 92.13 ± 1; PB_12_: 38.83 ± 1.28. Cells in L4 (%): PB_15_: 7.39 ± 0.66; PB_12_: 13.67 ± 1.24. Cells in L5 (%): PB_15_: 0.34 ± 0.28; PB_12_: 22.46 ± 3.45. Cells in L6 (%): PB_15_: 0.14 ± 0.14; PB_12_: 28.26 ± 2.87. AP_15→15_ and AP_12→12_ plots are reused from Fig. 1. A two-way ANOVA with post-hoc Tukey test was used.

**Figure S5b:** The layer position for all cells depicted in Figure 1 was plotted (N = recipient litter). In cortical areas with no anatomically distinguishable L4, the lowest third of L2/3 was arbitrarily considered as L4. Cells in L2/3 (%): AP_15→15_: 85.79 ± 2.58; AP_12→12_: 41.32 ± 5.71; AP_15→12_: 43.16 ± 3.63. Cells in L4 (%): AP_15→15_: 10.78 ± 2.24; AP_12→12_: 8.28 ± 3.18; AP_15→12_: 5.75 ± 1.54. Cells in L5 (%): AP_15→15_: 5.6 ± 2.16; AP_12→12_: 25.61 ± 3.99; AP_15→12_: 33.09 ± 2.15. Cells in L6 (%): AP_15→15_: 1.01± 0.79; AP_12→12_: 24.79 ± 4.07; AP_15→12_: 18 ± 3.90. A two-way ANOVA with post-hoc Tukey test was used.

**Figure S6a-c:** A subset of GFP^+^ neurons depicted in Fig. S4 was analyzed for molecular marker expression. Epor_PB15_: 380 cells from 5 pups (= N) were stained for CUX1 (% CUX1^+^: 94.96 ± 2.13); 380 cells from 5 pups were stained for CTIP2 (% CTIP2^+^: 0 ± 0); 304 cells from 3 pups were stained for TBR1 (% TBR1^+^: 1.89 ± 1.51). Epor_PB12_: 843 cells from 3 pups were stained for CUX1 (% CUX1^+^: 52.56 ± 6.1); 961 cells from 3 pups were stained for CTIP2 (% CTIP2^+^: 31.36 ± 1.29); 891 cells from 3 pups were stained for TBR1 (% TBR1^+^: 31.36 ± 1.29). The normalized radial position of labeled and non-labeled GFP^+^ neurons was plotted for 50 – 100 randomly selected cells for each condition and the total percentage of all cells expressing the respective marker were plotted. AP_15→15_ and AP_12→12_ plots and images showing expression of CUX1, CTIP2 and TBR1 within the neocortical layers are copied from Fig 2. A one-way ANOVA with post-hoc Tukey test was used.

**Figure S6d:** The fraction of EdU^+^ neurons at P7 from 7 donor litters across all conditions is shown.

**Figure S9a:** Three sections from different rostro-caudal levels from 3 pups (= N) were used to quantify the number of FT^+^ TBR2^+^ Ki67^+^ cells 10 hours after FT labeling at E15.5 (% FT^+^ TBR2^+^ Ki67^+^: 70.99 ± 2.81).

**Figure S9b:** The number of cells per integration site (oligoclone size) was analyzed pooling all cells from previous experiments. AP_15→12_: 8.11 ± 1.02; IP_15→12_: 3.77 ± 0.89.

**Figure S9d:** A subset of EdU^+^ cells depicted in Fig. 4d-f was analyzed for molecular marker expression. The normalized radial position of labeled and non-labeled cells and the total percentage of cells expressing the respective marker were plotted. A total of 15 cells from 2 independent experiments were stained for CUX1 (% CUX1^+^: 93.75 ± 6.25); 63 cells from 3 independent experiments were stained for CTIP2 (% CTIP2^+^: 0.83 ± 0.83). Images showing expression of CUX1 and CTIP2 within the neocortical layers are the same as in Fig. 2.

**Figure S9e:** The radial position of all RFP^+^ EdU^−^ neurons from subset of experiments shown in Fig. 4d-f (22 cells from 2 independent experiments) was measured and normalized to the cortical thickness.

### Electrophysiology and collection of cells for single-cell RNA sequencing

Three hundred µm thick coronal slices were prepared 24 hours after transplantation or FT labeling. Slices were kept for at least 30 minutes in artificial cerebrospinal fluid (aCSF) at 33°C (125 mM NaCl, 2.5 mM KCl, 1 mM MgCl_2_, 2.5 mM CaCl_2_, 1.25 mM NaH_2_PO_4_, 26 mM NaHCO_3_ and 11 mM glucose, saturated with 95% O_2_ and 5% CO_2_) before recording. The slices were then transferred in the recording chamber, submerged and continuously perfused with aCSF. The internal solution used for the experiments contained 140 mM potassium methansulfonate, 2 mM MgCl_2_, 4 mM NaCl 0.2 mM EGTA, 10 mM HEPES, 3 mM Na_2_ATP, 0.33 mM GTP and 5 mM creatine phosphate (pH 7.2, 295 mOsm).

For resting membrane potential recordings, immediately after the whole-cell configuration, resting membrane potential was measured in current-clamp mode. Membrane potential was monitored every 10 seconds and averaged for 6 consecutive acquisitions, within the first 2 minutes after the whole-cell configuration establishment. V_m_ remained stable within conditions and was significantly different across conditions throughout the duration of the recording, indicating that V_m_ recordings are not influenced by cytoplasmic dilution with the patch pipette solution. Recordings were amplified (Multiclamp 700, Axon Instruments), filtered at 5 kHz and digitalized at 20 kHz (National Instrument Board PCI-MIO-16E4, IGOR WaveMetrics), and stored on a personal computer for further analyses (IGOR PRO WaveMetrics). Values are represented as mean ± SEM.

For collection of cells for single-cell RNA sequencing, FT-labeled non-transplanted or RFP^+^ transplanted progenitors located in the ventricular zone were patched with a patch pipette containing 1 μl of internal solution supplemented with 1U/mL of RNase inhibitor (Takara). To facilitate the aspiration of the cell, low pipette resistance was used (4-3 MΩ). Once in whole cell configuration, a slit depression was applied to the pipette to aspirate the intracellular content. The complete aspiration of the cell was observed under high magnification (Nikon Eclipse FN1, 60x lens) as retraction of the cytoplasm and total aspiration of the nucleus. The patch pipette was then slowly retracted and the pipette tip containing the cell content was broken into a PCR RNase free Eppendorf containing 8 μl of lysis buffer and stored at −80°C until further processing.

### Single-cell RNA sequencing

cDNA synthesis and preamplification were performed using the SMART-Seq v4 3’ DE Kit following the manufacturer’s instructions (Takara). Single cell RNA-sequencing libraries of the cDNA were prepared using the Nextera XT DNA library prep kit (Illumina). Libraries were multiplexed and sequenced according to the manufacturer’s recommendations with paired-end reads using the HiSeq4000 plat-form (Illumina) with an expected depth of 1M reads per single cell, and a final mapping read length of 50 bp. Each pool contained cells from different collection days and conditions. All single cell RNA capture and sequencing experiments were performed within the Genomics Core Facility of the University of Geneva. The sequenced reads were aligned to the mouse genome (GRCm38) using STAR aligner. The number of reads per transcript was calculated with the open-source HTSeq Python library. All analyses were computed on the Vital-It cluster administered by the Swiss Institute of Bioinformatics.

### Single-cell RNA sequencing analysis

Cell filtering: A total of 91 cells (AP_12_: 31 cells, AP_15_: 31 cells, AP_15→12_: 29 cells) were sequenced. Cells expressing < 2000 genes or > 12% of mitochondrial genes were excluded (N = 26 cells). A total of 65 cells were analyzed (AP_12_: 20 cells, AP_15_: 26 cells, AP_15→12_: 29 cells: 19 cells). Gene expression was normalized to reads per million (rpm) and log transformed. Cell cycle effect was corrected for using the ccRemover package. We used the Support Vector Machine (SVM) method implemented in the bmrm R package to classify the age of progenitors with genes showing minimal expression (10,407) as previously described (Teo et al., 2010). The model was trained with a subset of AP_12_ and AP_15_ progenitors and the 30 most weighted genes were used for leave-one-out cross validation of additional AP_12_ and AP_15_ progenitors and prediction of AP_15→12_ cells. The same method was applied to build a model to classify AP_15_ and AP_15→12_, selecting for 100 most weighted genes.

---

**Figure S1:**
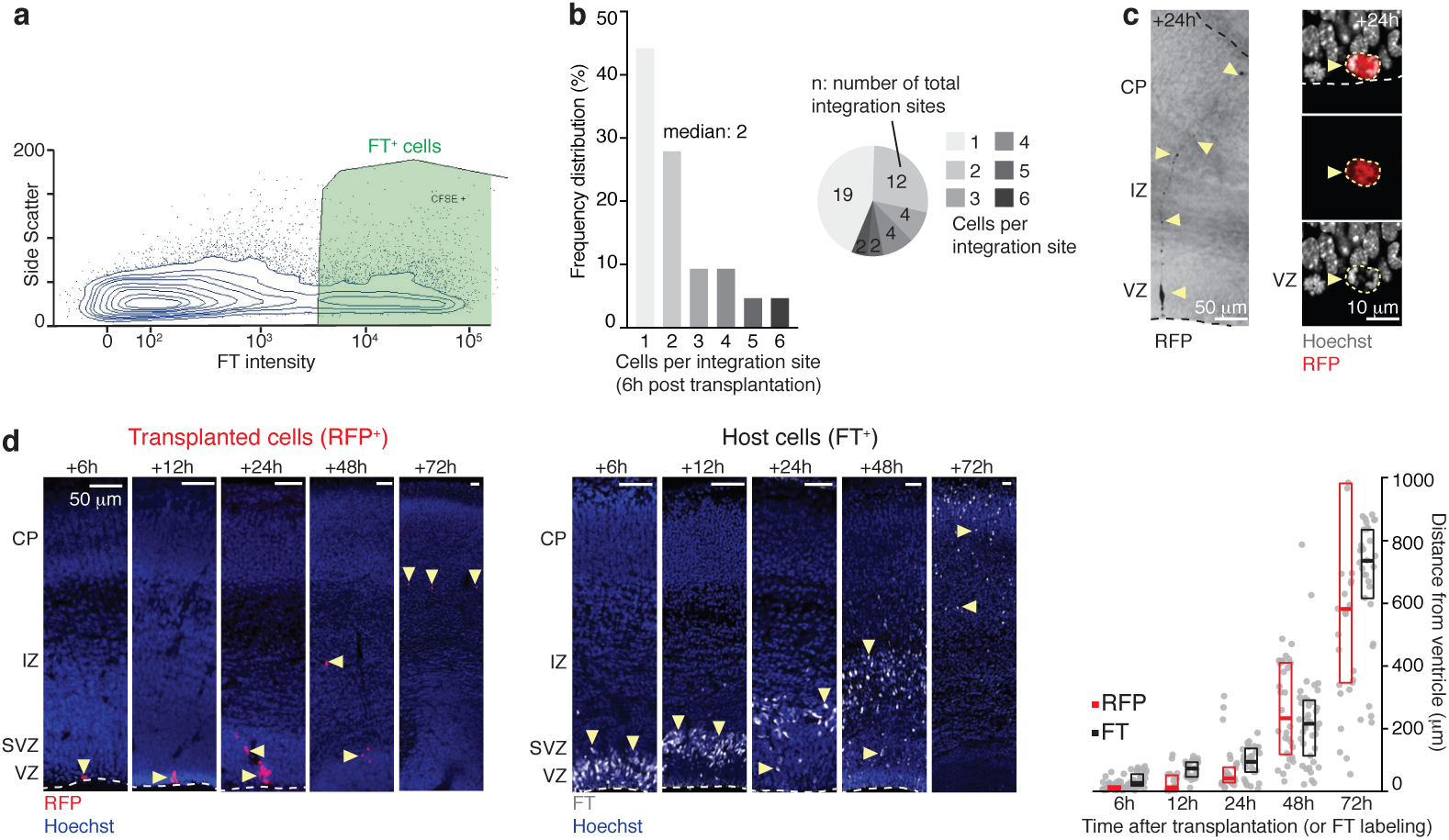
Donor APs rapidly integrate the VZ and behave normally. **a**, FAC-sorting of FT^+^ cells 1 hour after labeling. **b**, Donor APs integrate the host VZ at discrete sites. Only a few APs (median = 2) are present at each site, such that daughter neurons found at single integration sites at P7 are likely born from a small number of initial APs (“oligoclonal” analysis, see Fig.1g and Fig. 4b,f). **c**, Left: Photomicrograph of a transplanted AP showing a radial glia morphology. Right: Juxtaventricular mitosis in a transplanted AP. **d**, The progeny of transplanted APs progressively migrate towards the cortical plate. The time course of this migration is similar to that of the host cells, as assessed by comparison with the migration of FT-labelled endogenous cells. CP: cortical plate; FT: FlashTag; IZ: intermediate zone; SVZ: subventricular zone; VZ: ventricular zone.

**Figure S2:**
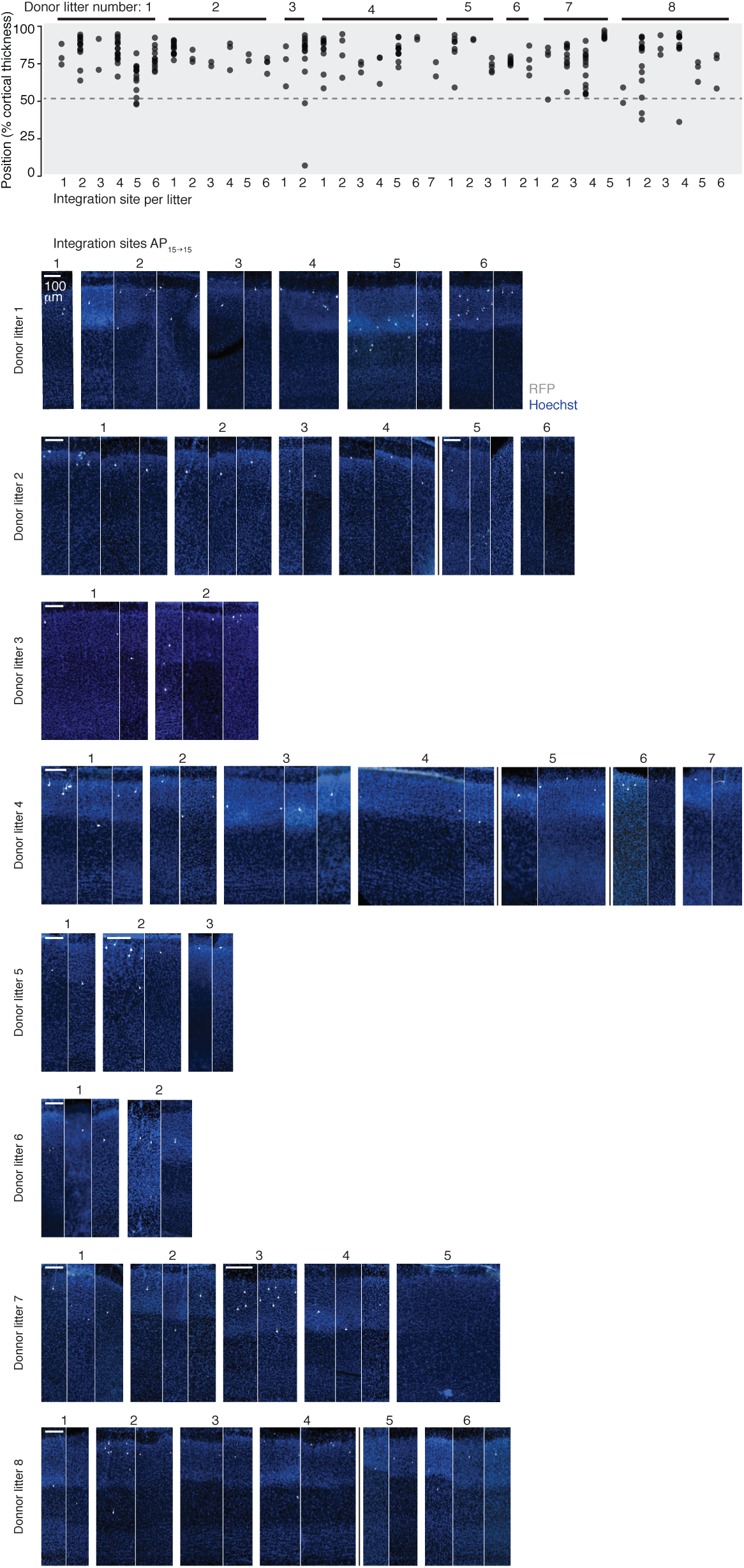
Analysis of single integration sites: AP_15→15_. Isochronically transplanted E15.5 APs (AP_15→15_) essentially generate SL neurons. Photomicrographs: within each donor litter, illustrations are clustered by integration site. When applicable, a vertical black line delineates distinct host pups within a given litter.

**Figure S3:**
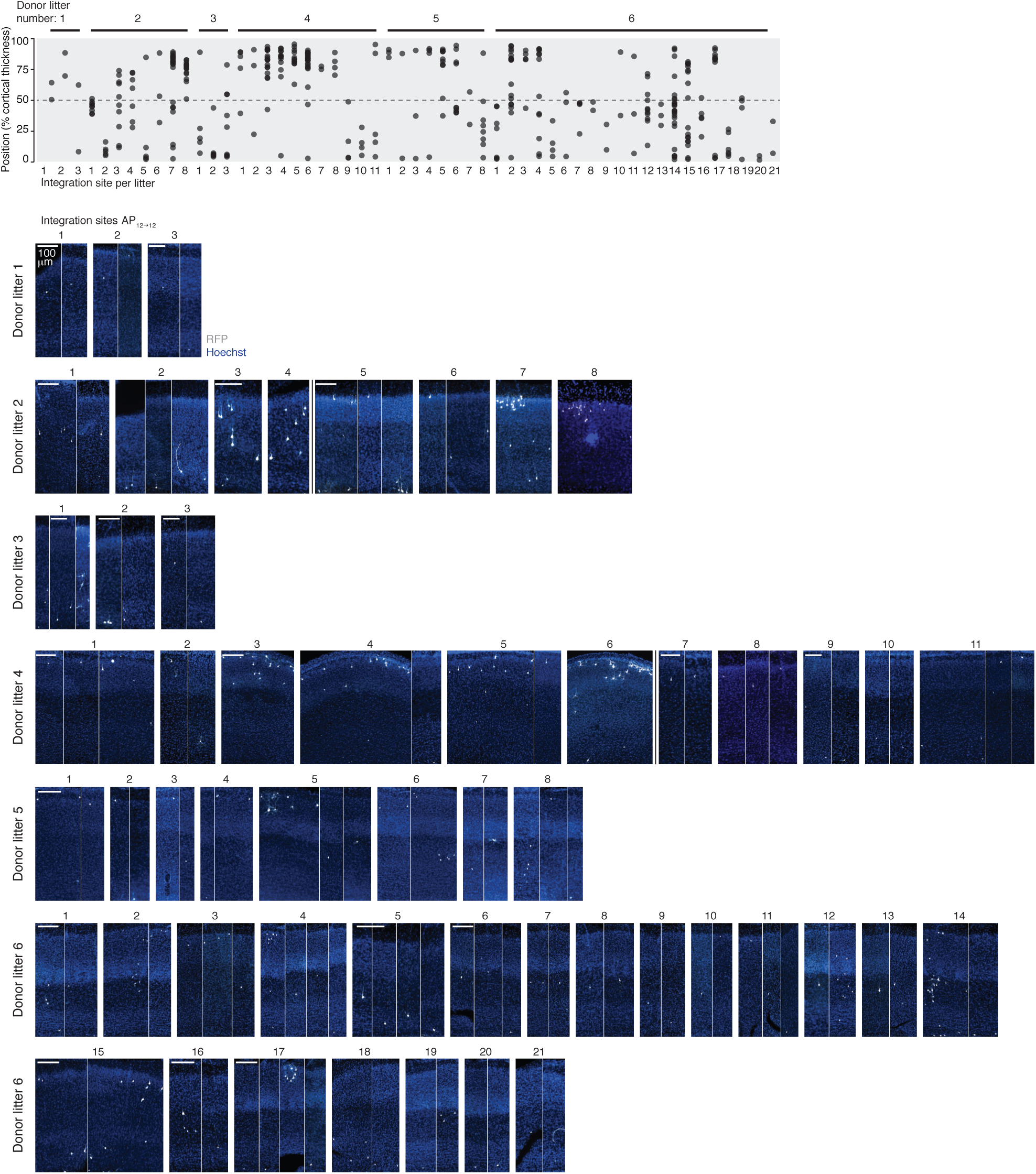
Analysis of single integration sites: AP_12→12_. Isochronically transplanted E12.5 APs (AP_12→12_) generate DL and SL neurons. Photomicrographs: within each donor litter, illustrations are clustered by integration site. When applicable, a vertical black line delineates distinct host pups within a given litter.

**Figure S4:**
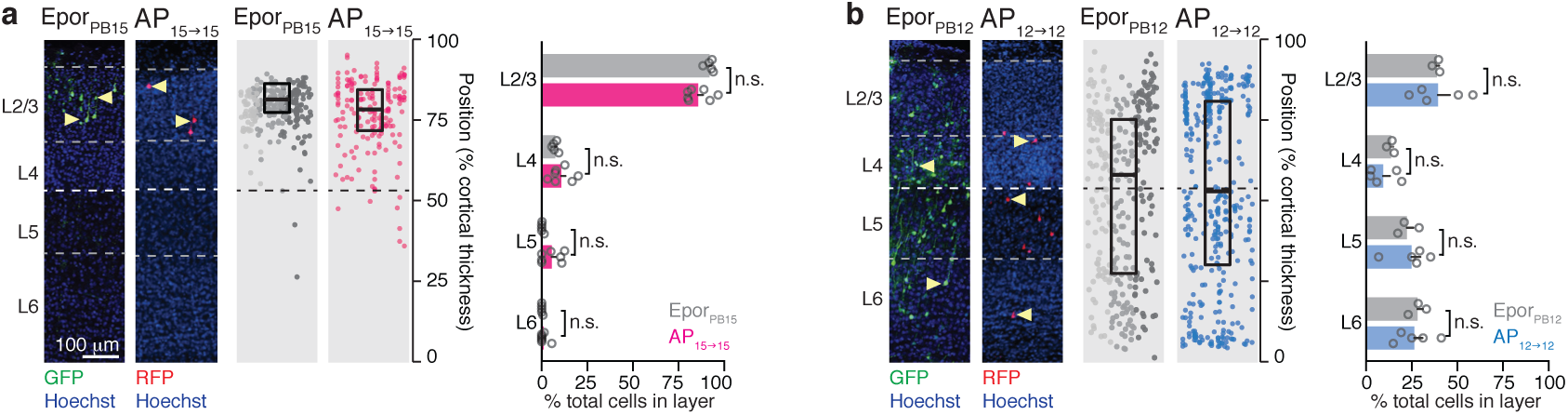
The transplantation procedure does not affect the neurogenic competence of APs. **a**, The laminar distribution of daughter neurons in the AP_15→15_ condition is replicated by *in utero* electroporation of a piggyBac-transposon construct at E15.5, in the absence of transplantation. **b**, The laminar distribution of daughter neurons in the AP_12→12_ condition is replicated by *in utero* electroporation of a piggyBac-transposon construct at E12.5. a-b: Two-way ANOVA with post-hoc Tukey test.

**Figure S5:**
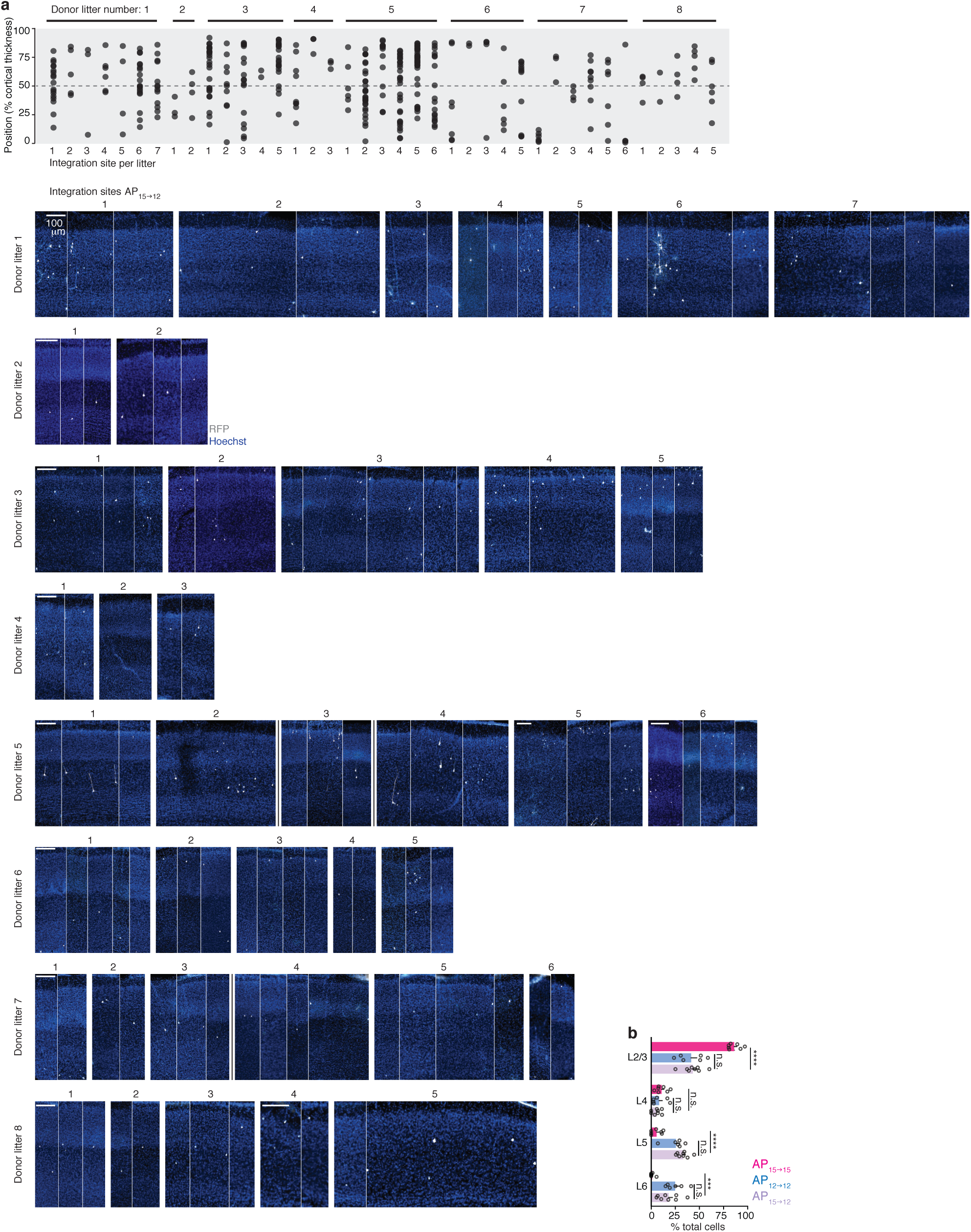
Analysis of single integration sites: AP_15→12_. **a**, E15.5 APs transplanted into an E12.5 host (AP_15→12_) generate DL and SL neurons. Photomicrographs: within each donor litter, illustrations are clustered by integration site. When applicable, a vertical black line delineates distinct host pups within a given litter. **b**, Laminar distribution of daughter neurons across conditions. AP_15→15_ and AP_12→12_ distribution plots are copied from Fig. S4 to allow for direct comparison across conditions. Two-way ANOVA with post-hoc Tukey test. ***: P < 0.001, ****: P < 0.0001.

**Figure S6:**
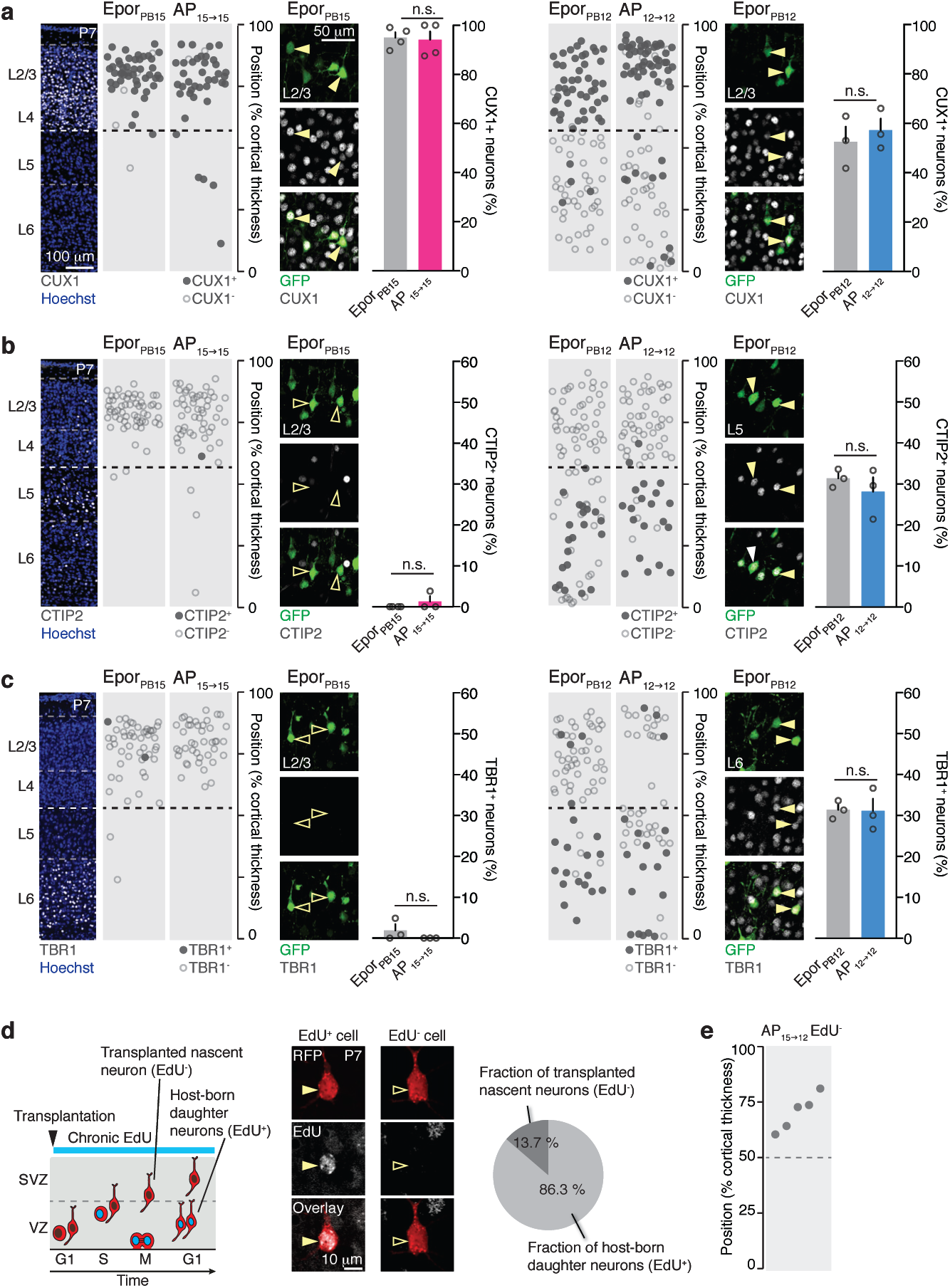
Molecular markers following piggybac electroporation; approach to identify transplanted nascent neurons. **a**, SL neurons in the isochronic transplantation and piggybac (PB) electroporation conditions both express CUX1. **b**, Daughter neurons of PB_15_ and AP_15→15_ do not express CTIP2. In PB_12_ and AP_12→12_ conditions, CTIP2^+^ daughter neurons are located in DL. **c**, Daughter neurons of PB_15_ and AP_15→15_ do not express TBR1. In PB_12_ and AP_12→12_ conditions, TBR1^+^ daughter neurons are located in DL. a-c: One-way ANOVA with post-hoc Tukey test. Photomicrographs showing pattern of expression of CUX1, CTIP2 and TBR1 are copied from Figure 2. **d**, Left: Schematic representation of the chronic EdU labeling approach used to distinguish between nascent donor neurons and neurons born in the host. Center: photomicrograph showing examples of an EdU^+^ and an EdU^−^donor neuron. Right: Quantification of the fraction of EdU^−^labeled neurons at P7 (*i.e.* transplanted cells which never underwent division in the host). **e**, Heterochronically transplanted E15.5 nascent neurons migrate to the superficial layers, as they would have done in their original host. M: mitosis; S: S-phase; SVZ: subventricular zone; VZ: ventricular zone.

**Figure S7:**
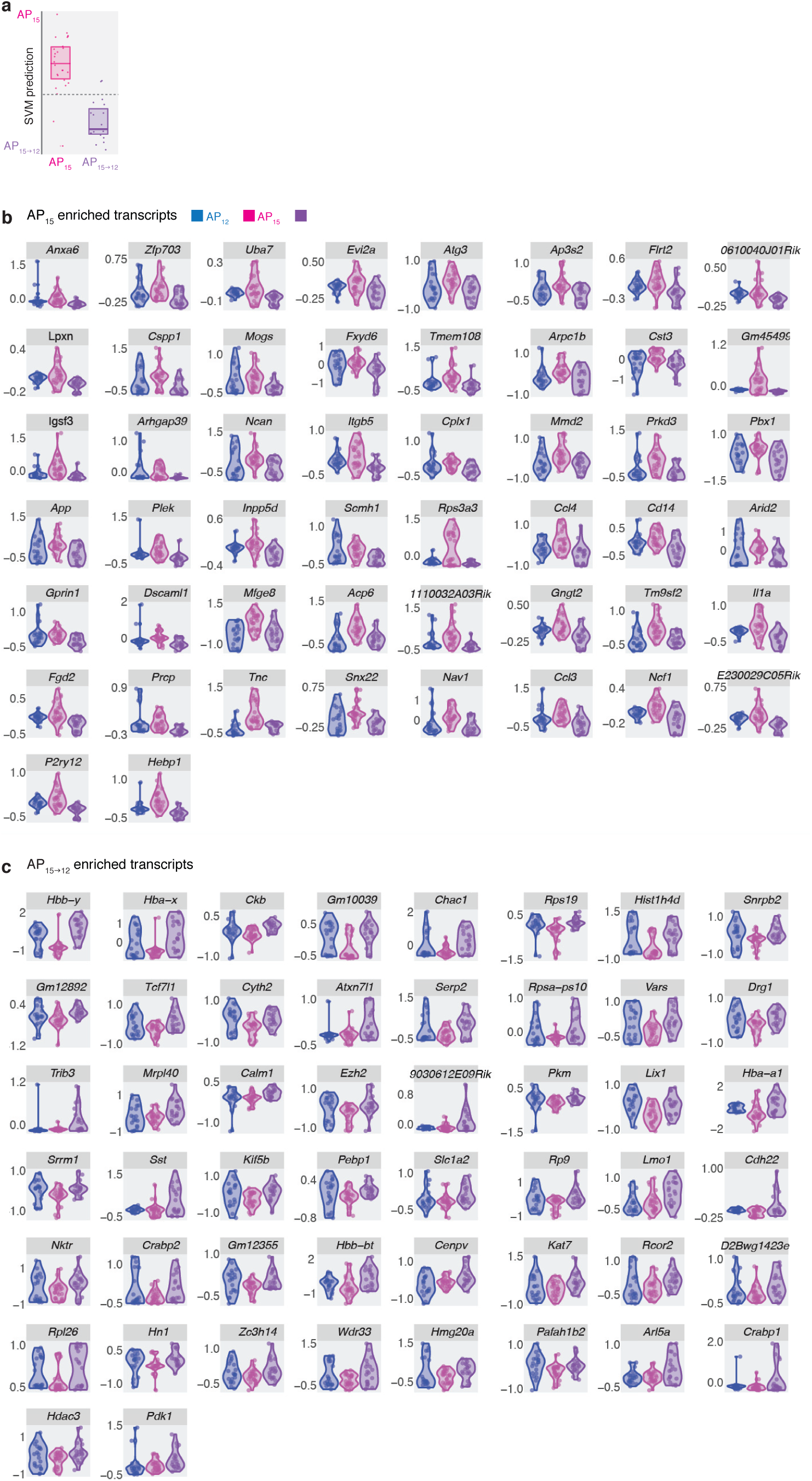
Repression of AP_15_-type transcriptional programs and re-induction of AP_12_-type transcriptional programs in AP_15→12_. **a,** SVM classification of AP_15→15_ and AP_15→12_. **b**, Expression of the AP_15_ transcripts used in the model. **c**, Expression of the AP_15→12_ transcripts used in the model.

**Figure S8:**
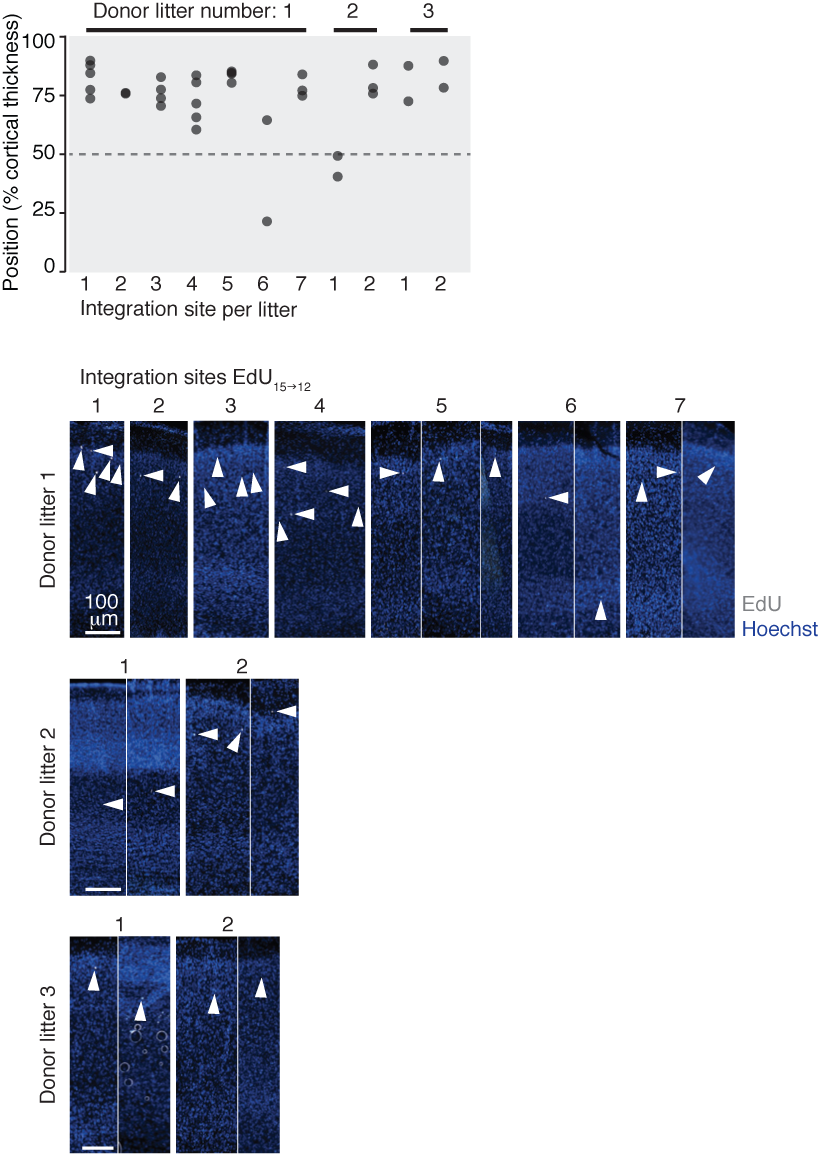
Oligoclonal analysis of single integration sites: EdU_15→12_. Heterochronically transplanted EdU-labeled progenitors (EdU_15→12_) essentially generate SL neurons. Photomicrographs: within each donor litter, illustrations are clustered by integration site.

**Figure S9:**
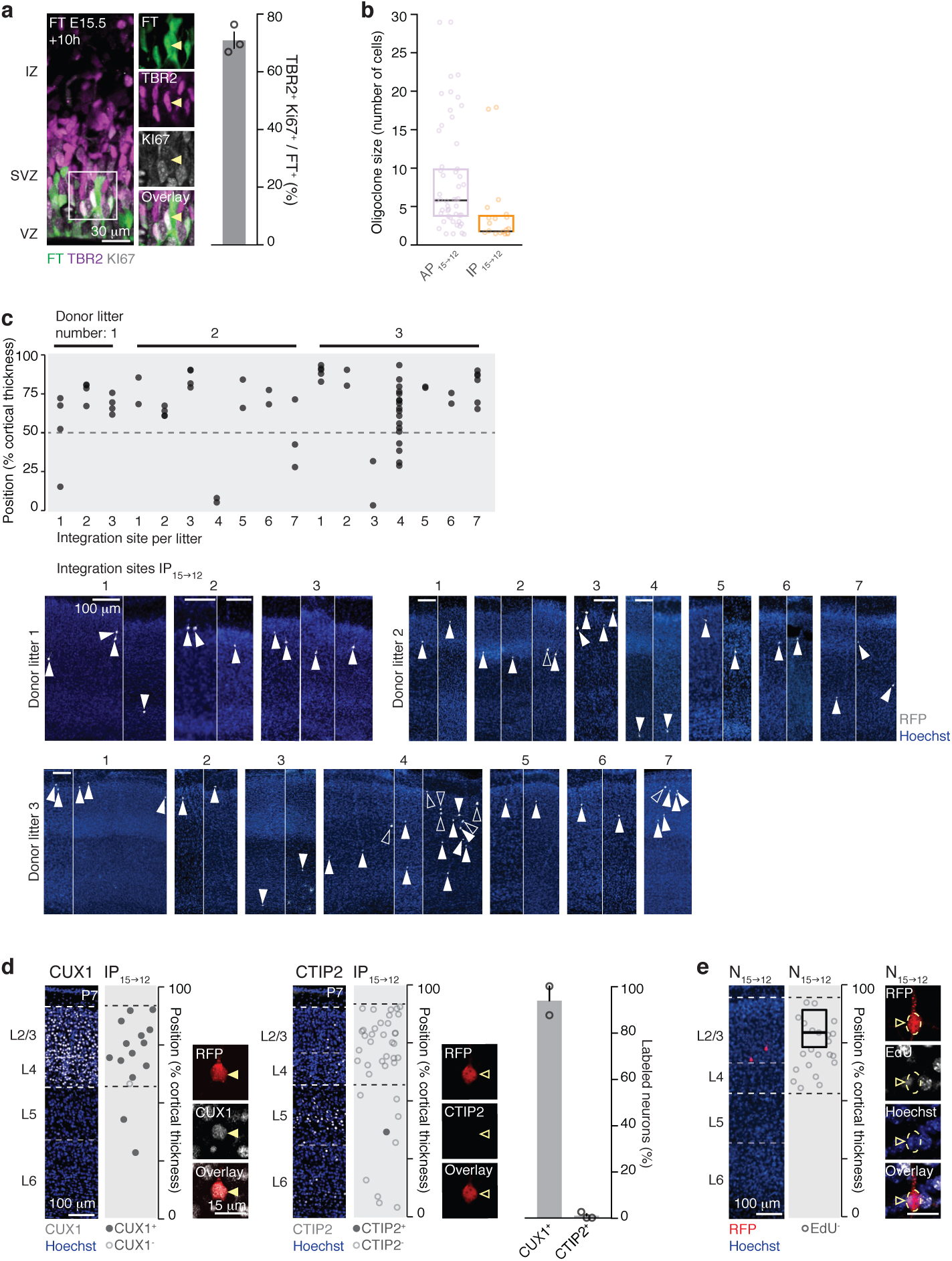
Heterochronically transplanted IPs (IP_15→12_) generate SL neurons. **a**, Ten hours after FT labeling, most cells have differentiated into IPs (*i.e.* KI67^+^ TBR2^+^ cells). **b**, Number of cells per integration site. Each point represents one oligoclone. **c**, IP_15→12_ essentially give rise to SL neurons. Photomicrographs: within each donor litter, illustrations are clustered by integration site. Only EdU^+^ neurons (filled arrowheads) were included in this analysis. **d**, IP_15→12_ daughter neurons express CUX1 but not CTIP2. **e**, Neurons that were post-mitotic at the time of transplantation migrate to SL. FT: FlashTag; IZ: intermediate zone; SVZ: subventricular zone; TBR2: IP marker; VZ: ventricular zone.

